# Movement trajectories as a window into the dynamics of emerging neural representations

**DOI:** 10.1101/2023.03.15.532848

**Authors:** Roger Koenig-Robert, Genevieve Quek, Tijl Grootswagers, Manuel Varlet

## Abstract

Transforming sensory inputs into meaningful neural representations is critical to adaptive behaviour in everyday environments. While non-invasive neuroimaging methods are the de-facto method for investigating neural representations, they remain expensive, not widely available, time-consuming, and restrictive in terms of the experimental conditions and participant populations they can be used with. Here we show that movement trajectories collected in online behavioural experiments can be used to measure the emergence and dynamics of neural representations with fine temporal resolution. By combining online computer mouse-tracking and publicly available neuroimaging (MEG and fMRI) data via Representational Similarity Analysis (RSA), we show that movement trajectories track the evolution of visual representations over time. We used a time constrained face/object categorization task on a previously published set of images containing human faces, illusory faces and objects to demonstrate that time-resolved representational structures derived from movement trajectories correlate with those derived from MEG, revealing the unfolding of category representations in comparable temporal detail (albeit delayed) to MEG. Furthermore, we show that movement-derived representational structures correlate with those derived from fMRI in most task-relevant brain areas, faces and objects selective areas in this proof of concept. Our results highlight the richness of movement trajectories and the power of the RSA framework to reveal and compare their information content, opening new avenues to better understand human perception.

## Introduction

The human brain’s astounding capacity for transforming sensory input into meaningful mental representations enables our adaptive behaviour in complex and continuously changing environments. While this capacity is now being increasingly investigated using neuroimaging, we show in this study that low-cost and widely available behavioural measures, human movement trajectories in particular, remain incredibly valuable to gain insight into the dynamics of emerging neural representations. Indeed, behavioural measures, such as reaction time and eye-tracking, have been for decades our main window into mental representations enabling gaining critical understanding of human cognition ^1–5^. The use of computer mouse-tracking movement trajectories is a more recent development in the behavioural toolbox ^6–9^. Mouse-tracking involves the continuous tracking of cursor trajectories towards one out of two or multiple choices, which has been found to be especially useful for measuring non-explicit processes such as self-control, emotion, ambivalence, moral and subliminal cognition^10–14^. Most importantly, movement trajectories have been proposed to inform not only about the end point of decisional processes, but also the temporal dynamics of decisions, revealing the emergence and duration of underlying neural representations ^6,9,15–17^.

However, the extent to which movement trajectories can index the continuous unfolding of cognitive processes, and more specifically, the transformation of visual inputs into meaningful neural representations, remains controversial ^18^. It is still highly debated whether movements, especially when performed under time constrains, can be modified by cognition once their execution has started. There are indeed studies suggesting that certain changes in trajectory might not be visually informed ^19^, that early visual perception might not be accessible by cognition ^20^, that the variability of movement outcomes might be mainly related to preparatory (pre-movement) neural activity ^21,22^, and that only single motor plans (i.e., a single choice, instead of competition among choices) would be represented in the motor cortex ^23^, thus challenging the hypothesis that the time-course of emerging neural representations can be captured via movement trajectories.

By combining movement trajectories and neuroimaging data, we show in this study that movement trajectories can provide a sensitive index of dynamic of neural representations. We show that observers’ mouse trajectories reveal the time course of decisional processes, capturing information about early visual representations and following their evolution (albeit delayed) towards their final stable state, instead of only reflecting the end product of decisional processes (i.e., a button press). We used publicly deposited neuroimaging data from Wardle et al. (2020) ^24^ which explored the time-course and brain areas supporting illusory face representations (face pareidolia). This phenomenon occurs when non-face stimuli elicit face perception due to their face-like visual features ^25,26^. Using images of human faces, pareidolic objects and non-pareidolic objects in combination with Magnetoencephalography (MEG) and functional Magnetic Resonance Imaging (fMRI), the aforementioned study revealed that illusory face representations emerge in earlier stages of visual processing, being resolved as objects later on. Using Representational Similarity Analysis (RSA), we compared these previously published neuroimaging data with mouse-tracking data we collected in an online face *vs*. object categorization task. We show that representational structures derived from movement trajectories matched those derived from MEG, following their temporal dynamics, albeit delayed. Furthermore, movement trajectories representational structures were found to be especially concordant with those derived from face and object selective brain areas as revealed by fMRI. Our results show that movement trajectories capture representational dynamics by reflecting individual stimuli differences, including their earlier visual processing stages, demonstrating decisive advantages over other behavioural measures focused on the end point of decisional processes only.

## Results

We recorded mouse trajectory data from a group of 77 online observers as they performed a face *vs*. object categorization task on the stimuli from Wardle et al. (2020) ^24^ (Figure 1A). To encourage participants to begin their classification movement early, each trial automatically terminated 800ms after stimulus presentation, or else when the participant clicked on a response box. Despite this time constraint, analysis of mouse trajectory endpoints showed that participants were highly accurate in categorising all three image categories (85.3, 80.68 and 82.36% for faces, pareidolic objects, and objects, respectively). Category information contained in trial-by-trial trajectories (see single trial examples in Figure 1B) was also reflected in conditional mean horizontal cursor position (Figure 1C): Trajectories corresponding to face and object images diverged from each other soonest (from 325ms), followed by those for faces and pareidolic objects (from 330ms). Trajectories corresponding to pareidolic objects and normal objects separated comparatively later (from 375ms) (p <.05; paired *t*-tests, FDR-corrected, q=0.05), with pareidolic object trajectories showing more attraction towards the face response box between 300-800ms (Figure 1C, inset). Note participants showed a slight initial bias towards responding ‘face’ (see Figure 1C&E), which could be caused by several factors (e.g., specialised face processing or treating it like a face vs not-face task), but this bias does not influence our analyses as we only examined relative position differences across exemplars.

**Figure 1.**
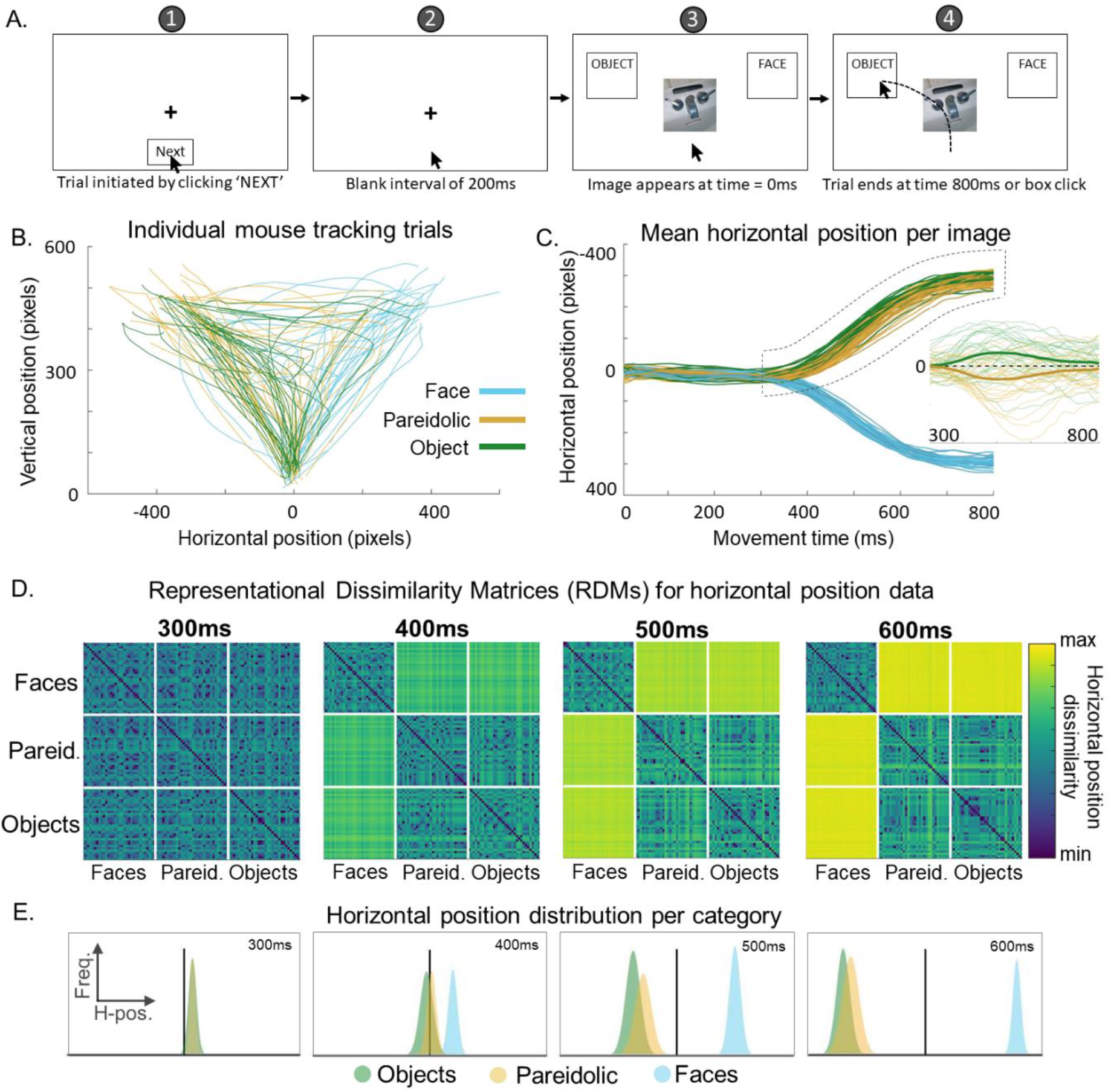
Mouse-tracking movement trajectory data. **A. Paradigm schematic**. We recorded online observers’ mouse trajectories in a face *vs*. object categorization task across 96 individual images (32 faces, 32 pareidolic objects and 32 matched objects) as used by Wardle et al. (2020) ^24^. We added a further 32 face images to equalise the probability of faces/objects, but did not include these additional face images for analysis. We instructed participants to only categorize human faces as faces, such that both pareidolic objects and normal objects had to be categorized as objects. (**1**) Each trial sequence began with a central fixation cross and a “Next” button that participants clicked on to initiate the trial. (**2**) When the trial commenced, participants first saw a 200ms blank interval to promote readiness for movement. (**3**) At time=0, either a face, pareidolic object or matched object appeared at fixation, along with two response boxes in the upper left and right corners of the screen. All images were presented once per block, and there were four blocks in total. The positions of the ‘OBJECT’ and ‘FACE’ response boxes swapped halfway through the experiment (i.e., after two blocks), with their initial positions counterbalanced across participants to guard against any right/left response biases. (**4**) Participants were instructed to move the cursor and click the appropriate response box as fast as possible, with each trial terminating 800ms after stimulus onset (or on box click). Both cursor landings and clicks on the correct box were considered as correct trials. Both correct and incorrect trials were included in subsequent analyses. **B. Individual mouse trajectories**. Individual mouse trajectory data for one block from a representative participant (96 trials). **C. Mean horizontal position over time for each exemplar in each category**. We took the horizontal component of the cursor movement (i.e., x-coordinate) as a time-resolved measure of the unfolding categorization response. For each image, we averaged x-position at each timepoint first within and then across participants (N=77), using these summary scores for further analyses. **Inset:** Image-wise deviation from the mean trajectory for objects and pareidolic objects. To visualise the distinction between object and pareidolic objects more clearly, we subtracted the grand mean from each image’s mean trajectory between 300 and 800ms (indicated by the dashed window in the main plot). Trajectories for objects and pareidolic objects separate in opposite directions, with pareidolic trajectories showing greater attraction towards the ‘FACE’ response box. Thick lines are the averaged mean-subtracted trajectories for each category. **D. Representational dissimilarity matrices (RDMs) for horizontal position data**. RDM for movement data illustrate the representational structure across tested images based on the categorization movement data. We constructed the RDM at each timepoint by taking the pairwise difference in pixels along the horizontal axis between the mean trajectory for each image. This resulted in a 96×96 matrix with 4186 unique pairwise combinations at each time point from 5-800ms after stimulus onset (step size = 5ms). RDMs at 300, 400, 500 and 600ms are shown for reference. Dissimilarities are shown as log2(distance) for display purposes. **E. Category distributions of horizontal position**. The distributions of horizontal positions for each object grouped by category are shown as histograms at 300, 400, 500 and 600ms. Faces started to separate from objects and pareidolic objects around 400ms, and remain separate over time. While the difference between pareidolic objects and objects was smaller than their difference to faces (given that both pareidolic objects and objects were categorized as objects), the distributions for pareidolic objects and objects remained offset, with the pareidolic object distribution indicating differences in their movement trajectory profiles and an attraction effect of faces over pareidolic.

Our primary goal was to examine the degree of representational overlap between our movement trajectory data and existing neuroimaging data for the same stimuli. We used Representational Similarity Analysis (RSA) ^27^ to abstract away from the native measurement units for these different datasets, projecting category distinctions reflected in MEG signals and horizontal position mouse trajectory data into the information domain via representational dissimilarity matrices (RDMs). Since the horizontal x-axis is the relevant dimension of categorization in our paradigm (i.e. go left for faces, go right for objects), the RDM series derived from time-resolved x-position data enables us to evaluate the emergence of category representations reflected in the unfolding movement trajectory (Figure 1D). We constructed these by calculating the pairwise trajectory distance along the x-axis (in pixels) between images for every time point (See Methods for details).

### Representational similarity between movement trajectories and MEG

In fusion analyses, high correlational values indicate shared representational structure across experimental measures, whereas low correlational values indicate rather different representational structures captured by each measure (Figure 2A). Time-time fusion analyses revealed that movement-derived representational geometries are comparable to those from MEG^24^ in both their structure and their ability to reflect category-specific visual processing among illusory faces, human faces and objects.

**Figure 2.**
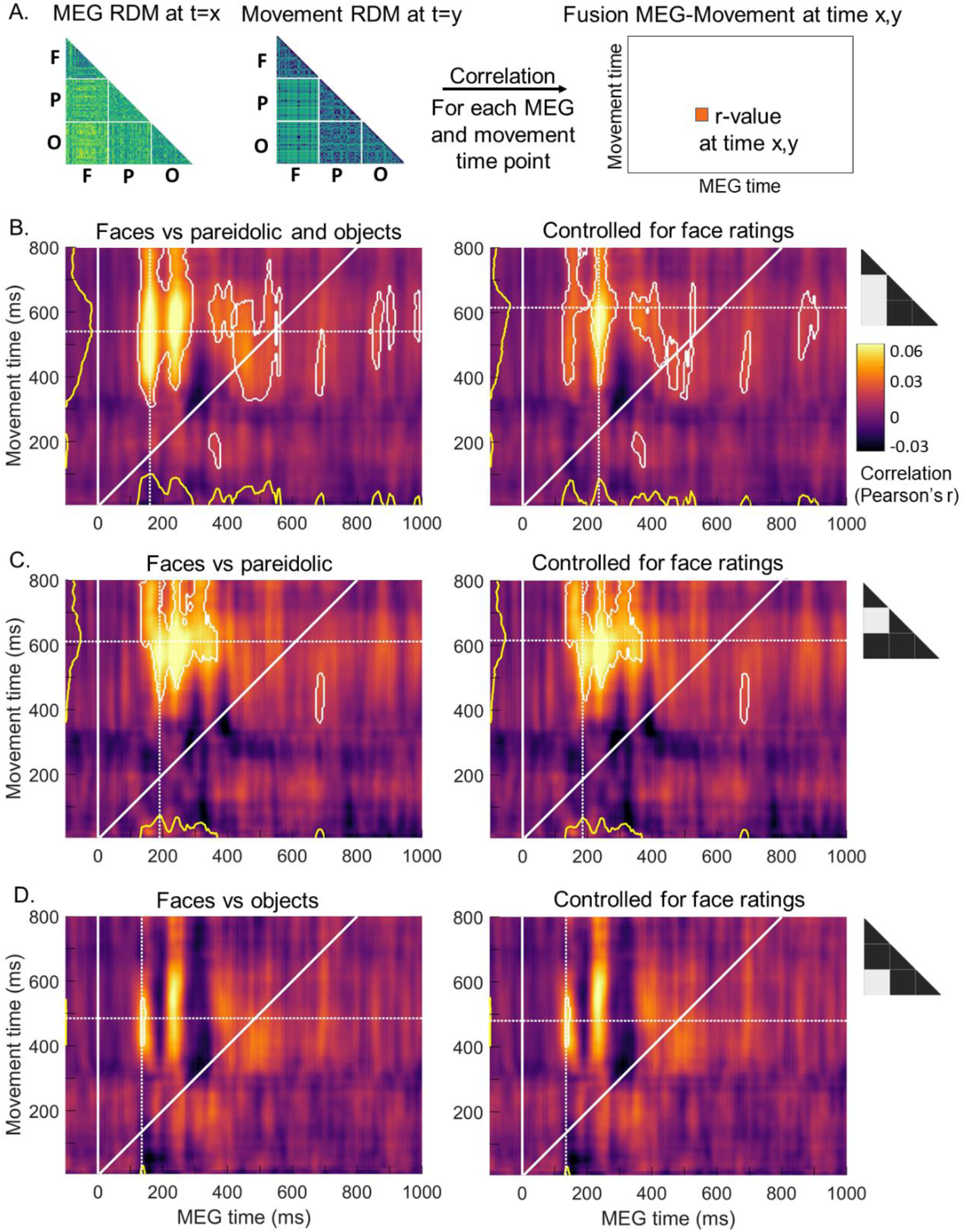
Representational overlap between movement trajectories and MEG responses. **A. MEG-movement fusion**. Fusion analysis evaluates the structural overlap between representations captured by different brain imaging techniques, behavioural measures and models ^28^. Here we compared the representational structures captured by MEG and movement data in a time-resolved manner to elucidate when these measures represent the stimuli (the 96 images dataset) similarly. RDMs from MEG data constructed by taking 1-correlation between the MEG activation patterns for each pair of stimuli (see Methods for details) were compared using Pearson’s correlation to RDMs constructed using movement data (Figure 1). Rather than computing correlations on the entire RDM, we selected parts of the RDM, thus focusing on the representational differences between faces and the rest of the stimuli as they hold information about the similarities and differences of the representations of faces, face-like and non-face objects. For each MEG and movement timepoint, we correlated a portion of the RDMs at each MEG and movement time combination. This produced time-time fusion maps with MEG time as the x-axis, movement time as the y-axis and the Pearson’s r-value colour-coded. We also calculated the partial correlations between MEG and movement data controlling for the variance explained by face ratings RDMs to check whether the representational similarities between the two could be accounted by simple face ratings (see ^24^ for details). **B. Faces *vs***. pareidolic objects and **objects**. Time-time MEG-movement fusions showed sustained common representational structures peaking at 160ms and 540ms for MEG and movement times respectively (as the maximum number of significant time-time points projected into each coordinate). Fusions controlled for face ratings showed a more restricted pattern of significant correlations peaking at 235ms and 615ms for MEG and movement data, indicating that some of the correlations between MEG and movement data are indeed explained by simple face ratings. **C. Faces *vs*. pareidolic**. Significant correlations were found to have a later peak than faces *vs*. pareidolic objects and objects, starting at 190ms and 610ms for MEG and movement times. Correlations controlled for face ratings showed virtually unchanged results compared to the non-controlled maps with significant correlations peaking at 185ms and 615ms for MEG and movement data, thus indicating that the face ratings do not capture the representational structure shared by movement and MEG for the face-pareidolic pairs. **D. Faces *vs*. objects**. While more constrained than faces *vs*. pareidolic objects and objects maps, faces *vs*. objects maps showed a peak at 135ms and 485ms of MEG and movement time, noticeably earlier than the faces *vs*. pareidolic subset. Fusion maps controlled by ratings showed virtually same results to the non-controlled maps, with significant correlations peaking at 135 and 480ms. White outlines represent significant correlations (one-sample t-test against 0, FDR-corrected for multi-comparisons q=0.05, cluster size threshold=50). Yellow lines represent the sum of significant time-time coordinates projected into each axis. Triangles represent the part of the RDMs selected (in light grey) for analyses.

To understand how the representations of human faces, pareidolic objects (or illusory faces, we use the two terms interchangeably), and objects reflected in movement data evolve over time, we focused our analyses on a subset of the RDMs that best represent the behavioural categorization task: i.e., faces *vs*. objects. In practice, this is achieved by selecting the RDM cells that represent dissimilarity between faces and pareidolic objects, and faces and normal objects (Figure 2B, light grey rectangles). Fusion analysis for this subset revealed clusters of significant correlation between MEG and movement representations that were shifted upwards from the diagonal (which represents identical movement and MEG times). This indicates that category representations reflected in movement trajectories lagged in time compared to those captured by MEG data. The peak of significant correlation between the two datasets (defined as the maximum number of significant points projected on each time axis) was located at 160ms and 540ms on the MEG and movement time axes, respectively.

Time-time fusion analysis for the RDM subset of faces vs. normal objects (Figure 2D) revealed that the robust neural distinction between faces and objects that arises very early in the MEG response (135ms) exhibited a sustained representational overlap with our mouse trajectory data between 400 to 550ms of movement time (peaking at 485ms). The same time-time fusion focused exclusively on the subset faces *vs*. pareidolic objects (Figure 2C) revealed shared representational structure across the two measures relatively later in time (peaking at 190ms and 610ms on MEG and movement times axis, respectively). The fact that movement-MEG representational overlap for this subset arises comparatively later (55 and 125ms in MEG and movement time, respectively) than for faces vs. objects is highly consistent with Wardle et al.’s ^24^ original report that maximal decoding arises later for faces *vs*. pareidolic objects (∼260ms) than for faces vs. normal objects (∼160ms).

### Movement trajectories *vs*. explicit ratings

Our results also showed that the information captured by movement trajectories go above and beyond the information captured by explicit ratings. We tested *how much* of the fusions between movement and MEG were explained by explicit face ratings’ RDM in the original paper ^24^. These face ratings (*face-likeness* of each image in a scale from 0 to 10) were completed by independent observers (N=20) in an online paradigm, see Methods and ^24^ for details. Face ratings indeed explained some of the correlations between movement and MEG representational structures, especially for the subset faces *vs*. pareidolic objects and objects, where controlled fusion maps showed more constrained regions of significant correlations (Figure 2B, right). For faces *vs*. pareidolic objects and faces *vs*. objects, fusion controlled maps remained virtually unchanged when compared to the original ones (Figure 2C-D, right), thus demonstrating that movement captured distinctly different representational information than face ratings.

### Representational similarity between movement trajectories and fMRI

Remarkably, fusion analysis with RDMs derived from fMRI data (Figure 3A) revealed that category representations reflected in movement data had structural overlap with those contained in face and object selective brain regions. We used representational structures obtained with fMRI from ^24^ in four category-selective brain areas: the fusiform face area (FFA), the occipital face area (OFA), the lateral occipital cortex (LO) and the parahippocampal place area (PPA) (Figure, 3B, see Methods for details on ROI definitions). Fusion analyses focused on faces *vs*. pareidolic objects and objects (Figure 3C) revealed that the representational geometry of faces, pareidolic, and object images as reflected in movement data were significantly correlated with geometries obtained in FFA (from 310ms of movement time), OFA (from 330ms) and LO (from 415ms), but not in PPA. These results are consistent with the selectivity of FFA, OFA, and LO brain areas for face and object perception and the role of PPA more oriented towards scene perception.

**Figure 3.**
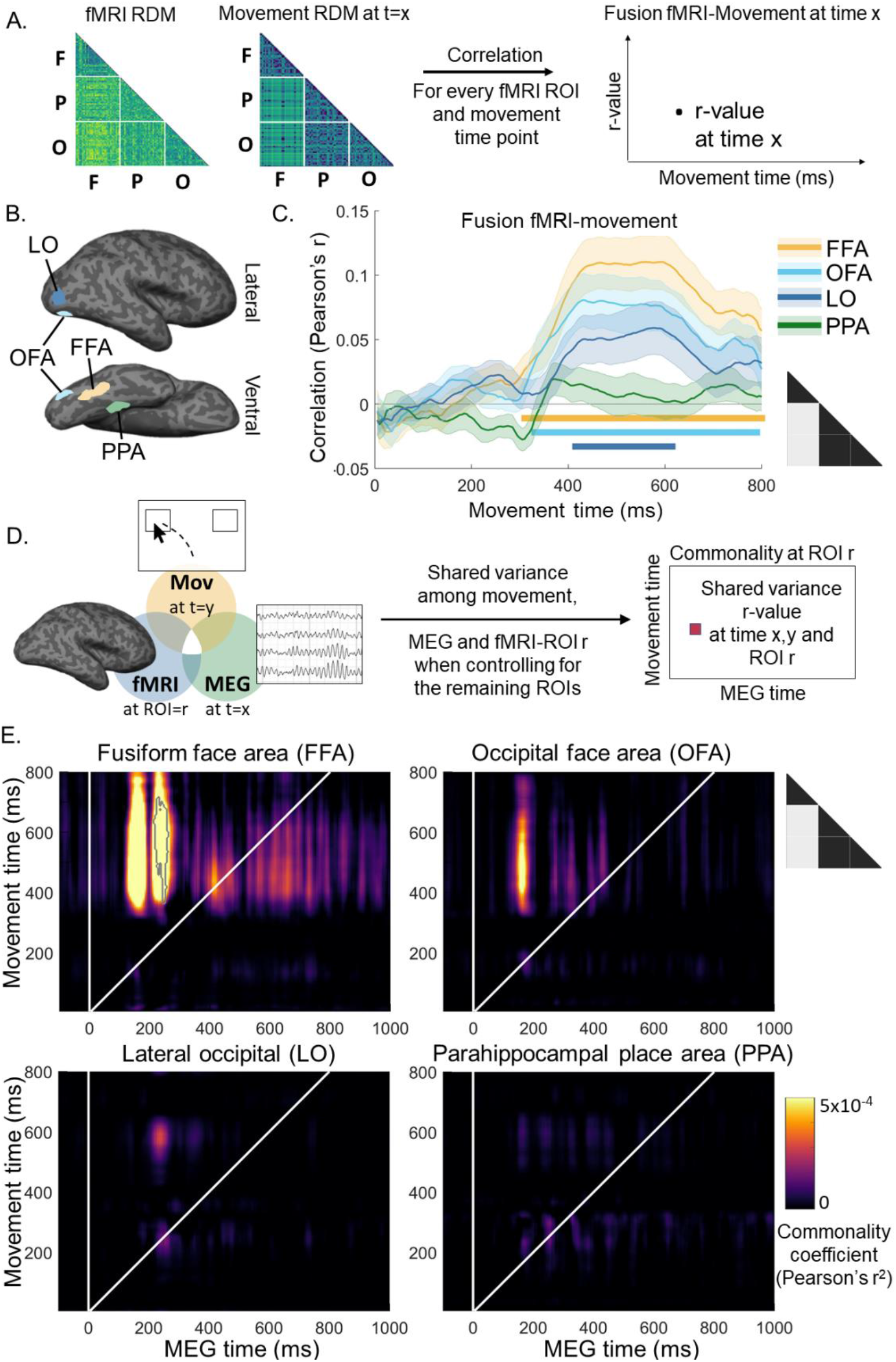
Representational overlap between movement trajectories and fMRI. **A. fMRI-movement fusion analysis**. We compared the representational structures captured by functional magnetic resonance imaging (fMRI) and movement in four ROI. RDM on each ROI were constructed by taking 1-correlation between the BOLD signal for each pair of stimuli (see Methods for details) and were correlated (Pearson) to RDMs constructed using movement data. **B. Category selective regions**. Four category selective regions were selected from fMRI recordings: the fusiform face area (FFA), the occipital face area (OFA), lateral occipital (LO) and parahippocampal place area (PPA). These ROIs were defined in an independent functional localizer experiment for each participant (see ^24^ for details). Two of these regions are selective to faces (FFA and OFA), one to objects (LO) and one to scenes (PPA), thus FFA, OFA and LO are expected to represent the differences between the stimuli set. Brains and ROI diagrams were modified from another study of our group and are shown for reference only. **C. Movement-fMRI fusions**. Rather than computing correlations on the entire RDM, we selected parts of the RDM to focus on the representational differences between faces and the rest of the stimuli as they hold information about the similarities and differences in coding for faces, face-like and non-face objects (light-grey on the black triangles). For each ROI and movement timepoint, we correlated a portion of the RDMs from both modalities to produce line plots representing the r-values as a function of movement time. **D. Movement-MEG-fMRI commonality analysis rationale**. We used a commonality analysis to investigate which brain areas shared representational structures with movement and MEG data. For each fMRI ROI, we calculated the partial correlation between MEG and movement RDMs when controlled by the variance in ROI=r minus the partial correlation of MEG and movement RDMs when controlled by the variance in all ROIs. **E. Time-time Movement-MEG-fMRI commonality maps**. The x and y axes correspond to MEG and movement time, respectively. The commonality coefficient (r^2^) is colour-coded. Commonality analyses showed that FFA significantly shared representational structures with MEG and movement. Note that commonality coefficients (r^2^) are often small in value (see for instance ^30,32,38^), as there are likely other sources of variance not accounted for in the models. Therefore, their statistical significance is often considered more important than their magnitude. Black outline represents significant correlations (one-sample one-tailed Wilcoxon signed rank test against 0, FDR-corrected for multi-comparisons q=0.05, cluster size threshold=50). Triangles represent the part of the RDMs selected (in light grey) for analyses.

The central contribution of FFA was further confirmed while investigating which brain regions shared common representations with movement and MEG using commonality analysis. Commonality analysis (Figure 3D) allows to identify the unique variance contribution of a single variable or predictor to the variance shared among multiple predictors ^29,30^. This analysis was used to test which brain areas from FFA, OFA, LO and PPA, contributed the most to the shared variance between movement, MEG and fMRI (see Methods for details). Commonality analyses revealed significant shared movement-MEG-fMRI representations in FFA, but not in other brain areas (Figure 3E). All in all, these results indicate that most of the shared variance between movement, MEG, and fMRI was explained by face-selectiveness in FFA, which is in line with its critical role in face recognition ^31,32^.

While commonality coefficients in OFA were not statistically significant, the latencies seen for FFA and OFA commonality maps could reveal temporal dynamics in the emergence of visual representations in these areas. The similar commonality latencies in OFA and the first responses in FFA could be interpreted as both areas producing face representations concurrently, thus challenging traditional posterior-to-anterior increase in visual hierarchy views ^33,34^, which have been contested by other studies ^35–37^.

## Discussion

Our results show that movement trajectories can be used to track the time course of unfolding neural representations, and that they capture representational structure beyond that reflected in behavioural measures focused on the end point of decisional processes (e.g., stimulus ratings). Our results also highlight the relevance of the Representational Similarity Analysis (RSA) framework to reveal the informational content in movement trajectories and compare movement data with other behavioural and neuroimaging data as well as theoretical models, which opens new avenues to understand human perception using the mouse-tracking paradigm.

### Movement trajectories as a window into emerging neural representations

Our results show that movement trajectories can be modified by cognition even under high time constraint after their execution has started. Moreover, classification movements in our study contained meaningful information about underlying neural representations of stimulus category. It is evidenced in this study by distinct early parts of movement trajectories for faces, illusory faces, and objects in our face *vs*. object categorization task consistent with their differences in early brain processing. Our results show that time-resolved representational structures derived from movement data were concordant with the unfolding of category representations measured by MEG as early as 120ms after stimulus presentation. Fusion analyses revealed a compelling overlap between stimulus representations reflected in movement and MEG data, with a notable offset between the two measures. The same representational structures evident in MEG data arise in trajectory data after a delay of some 380ms (e.g., for faces *vs*. pareidolic objects and objects). Importantly, fusion analysis with movement and fMRI data revealed that face, object, and pareidolic object representations derived from movement trajectory data show strong concordance with representations extracted from the BOLD response in the most task-relevant brain regions (i.e., regions with selectivity to faces and objects). Functional MRI-movement fusions indicated significant correlations between movement and FFA, OFA and LO, but not with PPA. These areas are selective to faces (FFA and OFA) and objects (LO) ^31,39^, in contrast to PPA that is involved in scene perception ^40^. The central role of FFA was further supported by the commonality analysis that showed that representational structures in FFA explained the shared information content between MEG and movement data better than any other brain area tested. Together, these results demonstrate the suitability of movement trajectories for measuring the time-course of emerging representations in the brain, including their early stages, which opens new possibilities for disentangling in future research stimulus features and processing stages driving human perception.

### Movement trajectories contain more information than explicit category ratings

Our results show that movement trajectories captured representational information that goes beyond that reflected in explicit category ratings. Indeed, when using face ratings from ^24^ to control the correlations between movement and MEG, we found that most of the similarities between movement and MEG representational structures were not explained by face ratings. This result further demonstrates the capacity of movement trajectories to capture time-course information about neural representations and their underlying intermediate representational categories. This is consistent with the ability of mouse-tracking to track non-explicit cognitive processes that otherwise are blurred (or resolved) by testing them explicitly, as in questionnaires or ratings ^10–14,41^. Our results support the assumption that hand movements are continuously updated by the dynamics of competing decisional processes ^42,43^, instead of representing their end product as explicit measures do. This property gives mouse-tracking the ability to reveal the dynamics of cognitive processes occurring in parallel and competing with each other. Our results also corroborate work linking neural representations to human reaction time behaviour ^44–48^and speak to the importance of linking neuroimaging data to behaviour ^49–51^. The rich information in movement trajectories may help reveal more subtle links like those between transient intermediate neural processing stages and early decision processes. Our mouse-tracking approach could be integrated in future neuroimaging studies to further explore how dynamic neural representations contribute to decisions.

### Representational similarity analysis framework to reveal information in movement trajectories

Our results underscore the utility of RSA as a powerful framework through which to marry informational content reflected in distinct behavioural and neuroimaging measures. RSA enables comparing information in movement trajectories with that in other systems, MEG, fMRI, and rating data in the present study. Combining mouse-tracking with neuroimaging through RSA has shown to be successful in previous studies to reveal the influence of specific brain areas in stereotypes ^52^, cultural-specific facial emotion and contextual associations ^53^ and social biases ^54^, but never before in a time-resolved manner as presented here. Time-resolved RSA enabled to test in this study similarities over time between the representational structures of movement trajectories and MEG data, and therefore, reveal the dynamics of neural representations developing after stimulus presentation. Movement trajectories combined with RSA offer endless possibilities to address new research questions as this unit-agnostic approach enables comparing information in movement trajectories with new theoretical models as well as increasingly available public EEG, MEG, fMRI, fNIRS, EMG and eye-tracking datasets. Remarkably, re-using public datasets does not necessarily imply asking the same research questions as the original study. Neuroimaging data from ^24^ used in the current study could be employed with new mouse-tracking tasks and/or participant populations to investigate for instance changes in the representation dynamics of faces, illusory faces and objects with face adaptation ^55^ and perceptual deficits ^56^.

### Conclusion

The flexibility and potential to answer a diverse range of questions makes the combination of mouse-tracking and publicly available neuroimage datasets through RSA a powerful choice for agile and accessible science. Mouse-tracking is as time and effort efficient as most explicit behavioural measures, while revealing more information, specifically the time-course of covert processes ^6,57^. Widely available and cost-effective, this method combined with RSA offers new opportunities to investigate the dynamical processes underlying human perception.

## Methods

### Images and neuroimaging data from Wardle et al 2020

We used the dataset from ^24^ consisting in 96 images (https://osf.io/9g4rz/). This dataset contained 32 human faces (faces), 32 illusory faces (pareidolic objects) and 32 matched objects (objects). For each illusory face image (pareidolic objects) a matched object image containing the same inanimate object(s) (although not pareidolic objects) was used, making these images comparable in their visual attributes. Human faces were also selected to reflect the high variance of the pareidolic objects and object images, containing different facial expressions, age, ethnicity, orientation and gender.

MEG, fMRI and explicit rating data was downloaded from the publicly available repository accompanying their publication at: https://static-content.springer.com/esm/art%3A10.1038%2Fs41467-020-18325-8/MediaObjects/41467_2020_18325_MOESM6_ESM.zip.

Briefly, MEG recordings from 22 participants were acquired using a 160-channel whole-head KIT MEG system. MEG data were down-sampled to 200Hz and PCA was applied for dimensionality reduction (retaining PCs explaining 99% of variance). MEG RDMs were constructed by taking 1-correlation (Spearman) between the MEG activation patterns for each pair of stimuli at each time point (N=221, from -100 to 1000ms after stimulus presentation). The MEG task consisted in the presentation (200ms) of the 96 visual stimuli (24 repeats of each stimulus). In each trial, images were tilted by 3° (left or right) and participants had to report the tilt direction.

Functional MRI recordings from 16 participants were acquired using a 3T Siemens Verio MRI scanner and a 32-channel head coil. A 2D T2*-weighted EPI acquisition sequence was used: TR= 2.5 s, TE= 32ms, FA= 80°, voxel size: 2.8 × 2.8 × 2.8 mm. The fMRI task was analogous to the MEG task with the difference that stimuli were presented for 300ms followed by a grey screen to complete a 4s trial. All stimuli were shown once per run and each participant completed 7 runs. Data were slice-time corrected and motion-corrected using AFNI. An independent functional localizer experiment using a different set of images was performed to define the category selective regions: FFA, OFA, LO and PPA. Functional MRI RDMs were builded by taking 1-correlation (Spearman) between the BOLD signal for each pair of stimuli (96×96) in each of the four category selective areas.

### Mouse-tracking participants

We tested first year students in psychology from Western Sydney University online through the SONA platform in exchange of course credits. Participants gave written informed consent to participate in the study, which was approved by the ethics committee of Western Sydney University. We tested 128 participants, from which, 109 participants completed the entire experiment. From these 109 participants, we discarded 17 participants as they had more than half of trials with no mouse tracking data (either because they chose not to move the mouse or because the data was unable to be collected by the browser). Further 15 participants were discarded as their performance was below 50% on the categorization task (possibly due to not performing the task). In total, datasets from 77 participants (68 females, age=24.4±0.9, righthanded=71, native English speakers=53) were considered for further analyses.

### Procedure

We used an online Web browser-based mouse-tracking face *vs*. object categorization paradigm. The experiment was built upon a publicly available code (https://github.com/mahiluthra/mousetracking_experiment) written in JavaScript using jsPsych 6 libraries ^58^ and hosted on Pavlovia ^59^. Experimenters had no direct interaction with the participants and the experiment ran locally in a web browser on participants own computer ^60^. The task started with a central fixation cross and a button marked “Next” at the bottom of the screen that the participant had to press before each trial, thus effectively repositioning the cursor at the bottom of the screen at the start of each trial (see Figure 1). After the “Next” button press, a blank screen with fixation cross was presented for 200ms in order to promote participant’s readiness to start moving. After the blank screen, an image of a human face (face), an object containing an illusory face (pareidolic objects), or a matched object (object) was shown at fixation. Two response boxes were presented in the upper left and right corners of the screen. One of them contained the word “OBJECT” and the other “FACE”. The position of the face and object response boxes was swapped halfway through the experiment (i.e., after two blocks), with the initial position of response boxes counterbalanced across to avoid right/left movement biases. Participants had 800ms to move the cursor to the response box to give the response to the categorization task. The trial ended after 800ms or when participants clicked on one of the boxes. In the mouse-tracking plugin, we set the recording of pointer position coordinates during the 800ms (or until button press) as fast as the local system could do (1ms) which effectively gave readings every 3 to 10ms, which were then linearly interpolated into 5ms temporal resolution. Correct trials were taken as those on which the participant clicked on the correct response box, and those where the cursor landed on the correct response box, even if there was no click. We presented the original 96 images used in ^24^: 32 human faces, 32 illusory faces (i.e., pareidolic objects), and 32 matched objects (see ^24^ for details). To avoid response biases due to a higher likelihood of objects compared to faces, we also included an additional 32 human faces which served to equalise the probability of objects and faces (additional face images not included in analyses). All 128 images appeared in each block; there were four blocks in total and participants could take a self-paced rest break in between each block as necessary.

### Mouse-tracking movement trajectory analysis

We analysed mouse-tracking data in MATLAB using in-house custom-developed scripts (https://osf.io/q3hbp/). We considered all trials for analyses (both correct and incorrect categorizations) as we hypothesized they jointly represent the unfolding of categorical representations. Empty values at the beginning of the mouse-tracking recordings (due to late onsets of the mouse movement) and the end (due to response box clicks before the 800ms deadline) were filled with NaN. We then linearly interpolated the data to 5ms intervals from 5 to 800ms. We then took mouse-tracking horizontal position (x-coordinate) as a time-resolved indicator of the categorization (and thus a time-resolved proxy of visual processing). Per each one of the 96 images, we averaged the horizontal position, first across trials within participants (4 trials per image), and then across participants (N=77). We thus considered 29568 individual mouse-tracking trials for analysis. These averaged responses were taken as a descriptor of the time-resolved face/object categorization of a given image. We then used a 20ms moving average window in order to smooth out the mouse-tracking movement trajectories. We used this same smoothing procedure (20ms moving average window) on the MEG data from ^24^.

### Representational similarity analysis (RSA) of movement data

Representational similarity analysis allows to compare different experimental measures by abstracting them in the information domain. The way an experimental measure (here pointer horizontal position) differs between two-given stimuli provides an estimation of how similarly (if the magnitudes match) or dissimilarly (if the magnitude difference is high) the two stimuli are represented ^27^. By calculating the differences between every pair of stimuli, the (dis)similarity matrix provides an estimation of how an experimental measure represents the whole experimental stimuli set. We constructed a representational (dis)similarity matrix (RDM) for each timepoint by calculating the absolute difference in horizontal position for every pair of images yielding 4560 unique pairs (excluding pairs of the same image). This produced 160 RDMs across the interval of 5 to 800ms after stimulus onset. RDMs organized from left to right and top to bottom with face images from positions 1 to 32, then pareidolic from 33 to 64 and then objects from 65 to 96. In order to focus on representational distinctions between specific categories, we then subset the RDMs in three different ways: 1) faces *vs*. pareidolic objects and normal objects (2048 unique pairwise comparisons), 2) faces *vs*. pareidolic objects (1024 unique pairwise comparisons), and 3) faces *vs*. normal objects (1024 unique pairwise comparisons).

### MEG-movement time-time fusion analysis

Fusion analyses allow to compare representational structures obtained from different experimental measures (for example, neuroimaging and behaviour) by correlating representational (dis)similarity matrices in a pair by pair basis ^27,28^. Since both MEG and movement data are time resolved, we compared RDMs from these two modalities at every combination of timepoints (35360 timepoints combinations). This temporal generalization approach (see ^61^ for a review) allowed us to identify delays in the onset of representational structures between modalities as well as sustained and repeated structures across time. We calculated the linear correlation (Pearson’s r) between RDMs from both modalities at every timepoint combination. For face-rating controlled maps, we calculated partial correlations (Pearson’s r) between MEG and movement RDMs while controlling for RDMs from face ratings.

### fMRI-movement fusion analysis

Similar to MEG-movement fusions, fMRI-movement fusion analyses were performed by comparing the representational structures from fMRI and movement via linear correlation (Pearson’s r). While fMRI data from ^24^ were not time resolved, there were 4 regions of interest (ROI) considered (FFA, OFA, LO and PPA). Correlations between RDMs for every fMRI ROI and movement timepoint were calculated to obtain a correlation value as a function of movement time.

### Movement-MEG-fMRI commonality analysis

In order to understand which brain areas from the four fMRI ROI shared information with the representational structures from the combination of movement and MEG, we used a commonality analysis ^29,30^. Commonality analysis allows to identify the unique variance contribution of a single variable or predictor to the variance shared among multiple predictors. This method has successfully been used in conjunction with RSA to compare how different predictors in the form of neuroimaging methods, models and tasks explain shared variance ^30,32,38,62^. Here, we focused on how 4 predictors, the fMRI ROI: FFA, OFA, LO and PPA, contributed to the shared variance between movement, MEG and fMRI. For each ROI (for example ROI1), we performed a commonality analysis by comparing the semi-partial correlations of all model variables except for the ROI whose contribution we wanted to isolate (Mov, MEG, ROI2, ROI3, ROI4), with the semi-partial correlation of all the model variables, including the selected ROI (Mov, MEG, ROI1, ROI2, ROI3, ROI4). We performed this analysis for each fMRI ROI as follows:

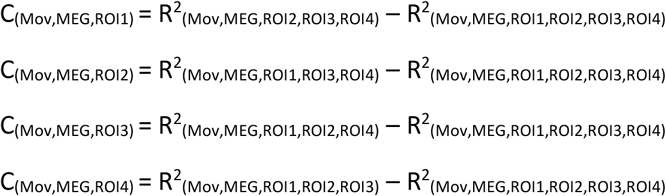

### Statistical inference

Time-time MEG-movement fusion maps and commonality maps’ correlations were tested via one-sample tests against 0 (h0: absence of correlation). We used two-sided t-tests for MEG-movement fusion maps and one-sided Wilcoxon signed rank tests for commonality maps across MEG participants (N=22). False discovery rate (FDR) ^63^ was used to control for multi-comparisons (type I errors or false-positives) with a q=0.05. Additionally, a cluster size of 50 time-time coordinates was set as the minimum size threshold for significance to avoid spurious results. Movement-fMRI fusions were tested using right-sided, one sample t-tests against 0 across fMRI participants (N=16) and multi-comparisons across movement timepoints were also controlled using false discovery rate (q=0.05).

## Data and code availability

Mouse-tracking data and MATLAB code to produce all results and figures are available at: https://osf.io/q3hbp/

